# Analysis of sub-kilobase chromatin topology reveals nano-scale regulatory interactions with variable dependence on cohesin and CTCF

**DOI:** 10.1101/2021.08.10.455796

**Authors:** Abrar Aljahani, Peng Hua, Magdalena A. Karpinska, Kimberly Quililan, James O.J. Davies, A. Marieke Oudelaar

## Abstract

Enhancers and promoters predominantly interact within large-scale topologically associating domains (TADs), which are formed by loop extrusion mediated by cohesin and CTCF. However, it is unclear whether complex chromatin structures exist at sub-kilobase-scale and to what extent fine-scale regulatory interactions depend on loop extrusion. To address these questions, we present an MNase-based chromosome conformation capture (3C) approach, which has enabled us to generate the most detailed local interaction data to date and precisely investigate the effects of cohesin and CTCF depletion on chromatin architecture. Our data reveal that *cis*-regulatory elements have distinct internal nano-scale structures, within which local insulation is dependent on CTCF, but which are independent of cohesin. In contrast, we find that depletion of cohesin causes a subtle reduction in longer-range enhancer-promoter interactions and that CTCF depletion can cause rewiring of regulatory contacts. Together, our data show that loop extrusion is not essential for enhancer-promoter interactions, but contributes to their robustness and specificity and to precise regulation of gene expression.

## MAIN

Mammalian gene expression patterns are controlled by enhancers, which form specific interactions with the promoters of their target genes to transfer activating signals^1^. Since these elements can be separated by large genomic distances, the specificity of enhancer-promoter interactions is dependent on the 3D organization of the genome^2^. Loop extrusion, mediated by cohesin and CTCF, organizes the genome into TADs^3–6^. Regulatory interactions between enhancers and promoters predominantly occur within TADs^7^ and disruption of TAD boundaries can result in ectopic enhancer-promoter interactions and changes in gene expression^3,8–16^.

It has been shown that depletion of cohesin, its loading factor NIPBL, and CTCF result in large-scale changes in the 3D organization of the genome and a loss of TADs^17–23^. However, surprisingly, the effects of these depletions on gene expression have been reported to be relatively mild. This poses an important question: to what extent are the specific regulatory interactions between enhancers and promoters that mediate gene expression dependent on loop extrusion?

To date, the effects of depletion of cohesin and CTCF on 3D chromatin structure have been analyzed by genome-wide Chromosome Conformation Capture (3C)^17–22,24^ and imaging approaches^21,25–27^. Although these studies have given important insights into the roles of cohesin and CTCF in organizing the genome, the resolution and sensitivity of the approaches used in these studies is not sufficient to characterize potential changes in sub-kilobase-scale chromatin architecture and enhancer-promoter interactions upon cohesin and CTCF depletion.

Conventional 3C methods^28,29^ are limited in their resolution by sequencing depth, library complexity and the distribution of the recognition sites of the restriction enzymes used for chromatin digestion^30^. The latter can be overcome by the use of MNase, which digests the genome largely independent of DNA sequence. MNase digestion was initially implemented in the Micro-C approach in yeast^31,32^. Micro-C has recently been adapted for use in mammalian genomes and has demonstrated a better signal-to-noise ratio compared to Hi-C and potential to generate contact matrices at high resolution^33–35^.

We have recently developed a novel MNase-based approach called Micro-Capture-C (MCC), which is capable of generating base pair resolution 3C data in mammalian cells^36^. This technique produces interaction profiles for selected viewpoints, but does not give an overview of the complete 3D structure of regions of interest. We have therefore extended the MCC approach and combined it with enrichment panels consisting of capture oligonucleotides tiled across genomic regions of interest (Figure 1a). This Tiled-MCC approach has a number of advantages over Micro-C, which enable the generation of data at higher resolution. These include maintaining cellular and nuclear architecture by using digitonin instead of traditional detergents; very deep data generation through high library complexity and targeted sequencing; and direct identification of ligation junctions. The Tiled-MCC approach has enabled us to generate the deepest local 3C matrices to date in mouse embryonic stem (mES) cells with high reproducibility (Supplementary Figure 1). Direct comparison of Tiled-MCC and Micro-C^34^ data clearly shows that Tiled-MCC is able to detect enhancer-promoter interactions and long-range contacts between CTCF-binding elements which are not detected by Micro-C (Figure 1b-d).

**Figure 1.**
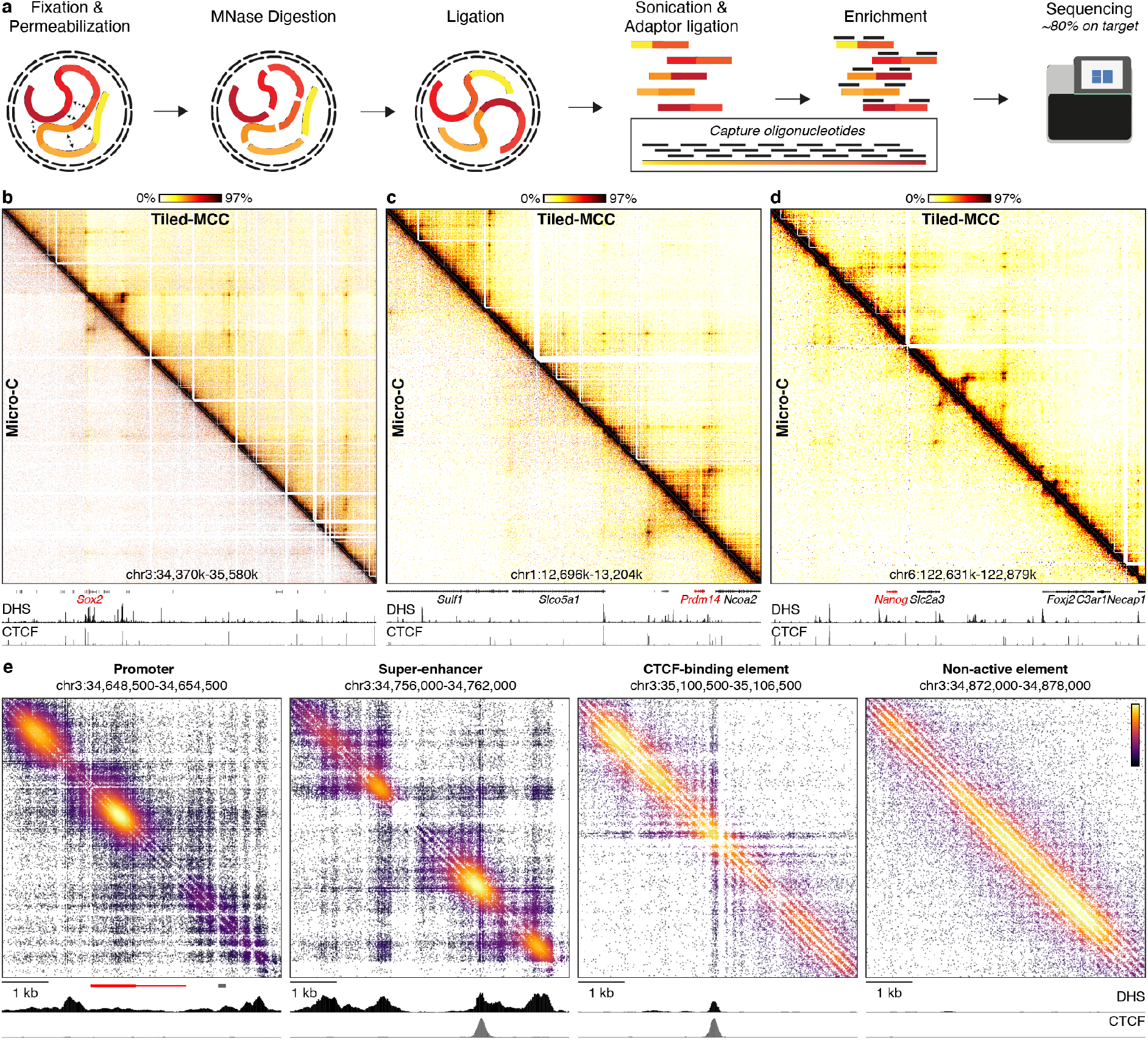
Tiled-MCC generates local contact matrices with substantially improved sensitivity and resolution. **a.** Overview of the Tiled-MCC procedure. Cells are initially cross-linked with formaldehyde and permeabilized with digitonin. The chromatin is subsequently digested with MNase, which is followed by proximity ligation. After DNA extraction, the MCC libraries are sonicated and ligated with indexed sequencing adaptors. Multiplexed libraries are subsequently enriched for regions of interest using panels of biotinylated capture oligonucleotides, optimized for hybridization to MNase-digested 3C libraries. Since MNase does not cut at specific sites in the genome, the panels are designed to densely cover regions of interest with oligonucleotides of 70 nucleotides in length and an overlap of 35 nucleotides. This optimized capture oligonucleotide design results in ~80% of reads on target. **b-d.** Comparison of Tiled-MCC (top-right) and Micro-C (bottom-left) contact matrices at 500 bp resolution of the *Sox2* (**b**), *Prdm14* (**c**), and *Nanog* (**d**) loci in mES cells. Gene annotation (genes of interest in red, coding genes in black, non-coding genes in grey), DNase hypersensitive sites (DHS), and ChIP-seq data for CTCF are shown below the matrices. The axes of the DHS and CTCF ChIP-seq profiles are scaled to signal and have the following ranges; DHS: *Sox2* = 0-4.46, *Prdm14* = 0-6.45, *Nanog* = 0-10.25; CTCF: *Sox2* = 0-1833, *Prdm14* = 0-2168, *Nanog* = 0-3092. **e.** Density plots of ligation junctions identified by Tiled-MCC showing the micro-topology of *cis*-regulatory elements in the *Sox2* locus. Gene annotation, DNase hypersensitive sites (DHS), and ChIP-seq data for CTCF are shown below the plots. The axes of the DHS and CTCF ChIP-seq profiles are fixed and have the following ranges: DHS = 0-5; CTCF = 0-1500.

Since Tiled-MCC libraries are sonicated to an average size of ~200 base pairs and sequenced with paired-end reads of 150 base pairs each, ligation junctions are sequenced directly and can be precisely reconstructed bioinformatically. This further increases the signal-to-noise ratio of the Tiled-MCC contact matrices and allows for generation of density plots of ligation junctions in which local chromatin structures can be analyzed with unprecedented resolution. These density plots reveal characteristic micro-topologies at *cis*-regulatory elements (Figure 1e, Supplementary Figure 2). Promoters and super-enhancers form complex nano-scale structures, characterized by sub-kilobase compartments of interaction. CTCF-binding elements form distinct patterns of phased nucleosomes and localized stripes. Interestingly, CTCF-binding sites within super-enhancers create localized regions of insulation between the different micro-compartments. Non-active chromatin is characterized by regular nucleosome structures without specific phasing patterns.

The advantages of Tiled-MCC make it a unique approach to investigate changes in local chromatin structure in detail upon depletion of architectural proteins. To study the role of cohesin and CTCF in mediating large- and fine-scale chromatin structure, we used mES cell lines in which the cohesin subunit RAD21 (SCC1) or CTCF can be rapidly degraded via an auxin-inducible degron (AID). To examine enhancer-promoter interactions in the absence of cohesin, we produced Tiled-MCC matrices of four well-characterized gene loci in RAD21-AID mES cells. We initially focused our analyses on the region containing the *Sox2* gene and its super-enhancer^37^. Consistent with previous reports^18,20^, cohesin depletion results in a near-complete loss of TAD structure and long-range interactions between CTCF-binding sites (Figure 2a, Supplementary Figure 3). A close-up view reveals that interactions between the *Sox2* promoter and its super-enhancer remain partially intact, despite loss of the interactions between the CTCF-binding sites surrounding these elements (Figure 2b). However, the interactions between the enhancers and promoter are reduced in intensity. This reduction in enhancer-promoter contacts corresponds to a small, but significant decrease in *Sox2* expression (Figure 2c). We next investigated the *Prdm14*, *Nanog* and *Pou5f1* loci, which are also regulated by well-characterized super-enhancers. In each of these loci we find a subtle decrease in enhancer-promoter interactions and gene expression following cohesin depletion (Supplementary Figure 4).

**Figure 2.**
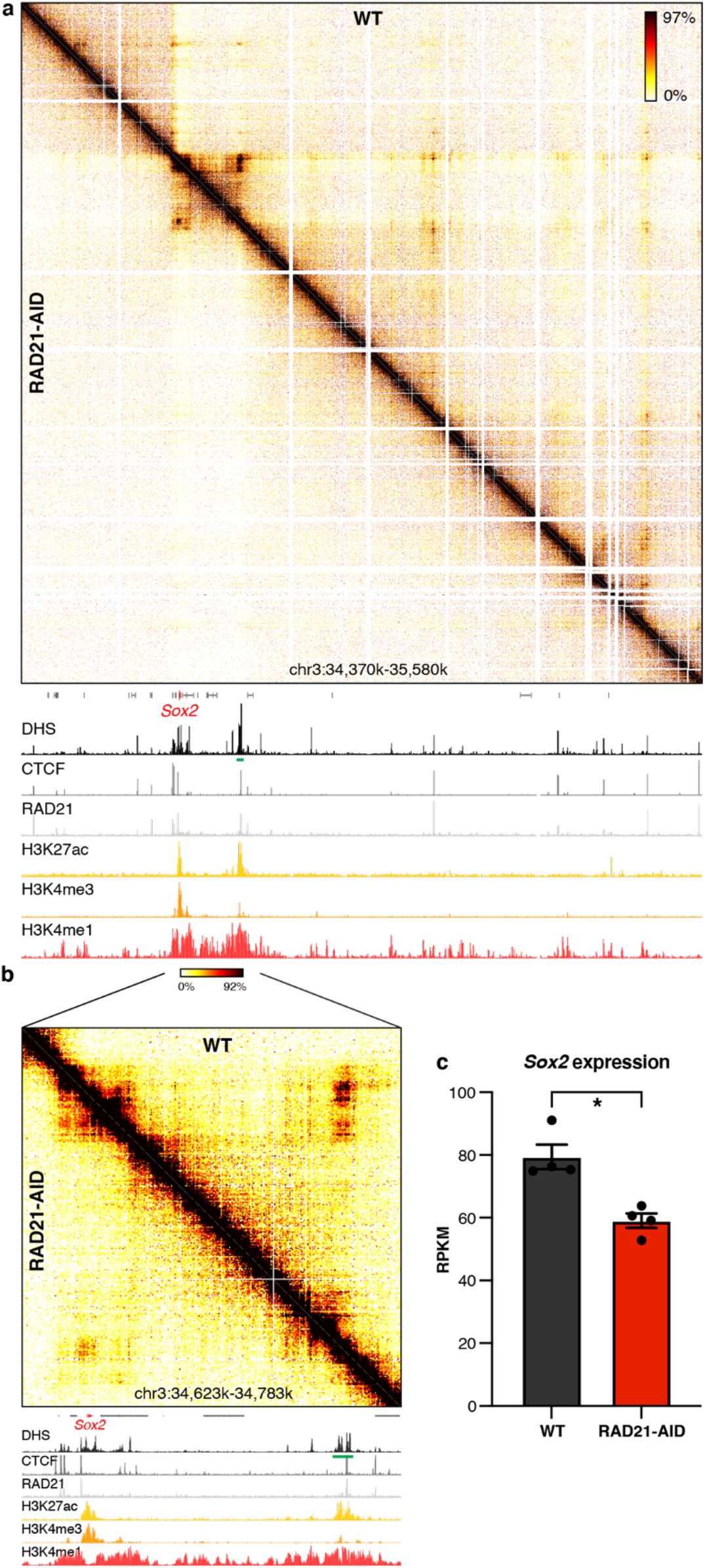
Cohesin depletion results in reduced enhancer-promoter interactions in the *Sox2* locus. **a.** Tiled-MCC contact matrices of the *Sox2* locus in wild type (WT) mES cells (top-right) and auxin-treated RAD21-AID mES cells (bottom-left) at 500 bp resolution. Gene annotation (*Sox2* in red, coding genes in black, non-coding genes in grey), DNase hypersensitive sites (DHS) and ChIP-seq data for CTCF, Cohesin (RAD21), H3K27ac, H3K4me3 and H3K4me1 are shown below the matrices. The axes of the DHS and ChIP-seq profiles are scaled to signal and have the following ranges: DHS = 0-4.46; CTCF = 0-1833; RAD21 = 0-3318; H3K27ac = 0-48; H3K4me3 = 0-82; H3K4me1 = 0-1826. Enhancers of interest are indicated in green below the DHS profile. **b.** Zoomed view of the data presented in panel a to highlight the interactions between the *Sox2* promoter and its enhancers. The axes of the DHS and ChIP-seq profiles are scaled to signal and have the same ranges as in panel a, except for the CTCF ChIP-seq profile, which is scaled 0-300. **c.** Expression of *Sox2* in untreated (left) and auxin-treated (right) RAD21-AID mES cells, derived from RNA-seq data, normalized for reads per kilobase of transcript, per million mapped reads (RPKM). The bars represent the average of n=4 replicates and the error bars indicate the standard error of the mean. * P = 4.46E-07.

Together, these results indicate that cohesin contributes to enhancer-promoter interactions. However, it is not clear if the effects of cohesin depletion are due to loss of TAD insulation or result from loss of active loop extrusion. To get further insight into the underlying mechanism, we studied the effects of CTCF depletion, which leads to a loss of TAD boundaries, but does not interfere with loop extrusion otherwise. Consistent with the literature^19,20^, Tiled-MCC data in CTCF-AID mES cells show that CTCF depletion results in a loss of TADs and interactions between CTCF-binding sites (Figure 3, Supplementary Figure 5). Furthermore, at the *Nanog* locus, we find that loss of CTCF leads to rewiring of the interactions between enhancers and promoters (Figure 3). Upon CTCF depletion, the enhancers form stronger contacts with the *cis-*regulatory elements of the downstream *Foxj2* gene, which leads to upregulation of its expression. We also observe the formation of ectopic enhancer-promoter interactions at the *Prdm14* locus (Supplementary Figure 6), which are consistent with previously reported effects of CTCF-binding site deletions in this locus^8,38^. However, CTCF depletion at the *Sox2* and *Oct4* loci does not result in changes in enhancer-promoter interactions or gene expression in mES cells (Supplementary Figure 6). These results show that CTCF is important for the specificity of enhancer-promoter interactions, but not required for their formation or maintenance.

**Figure 3.**
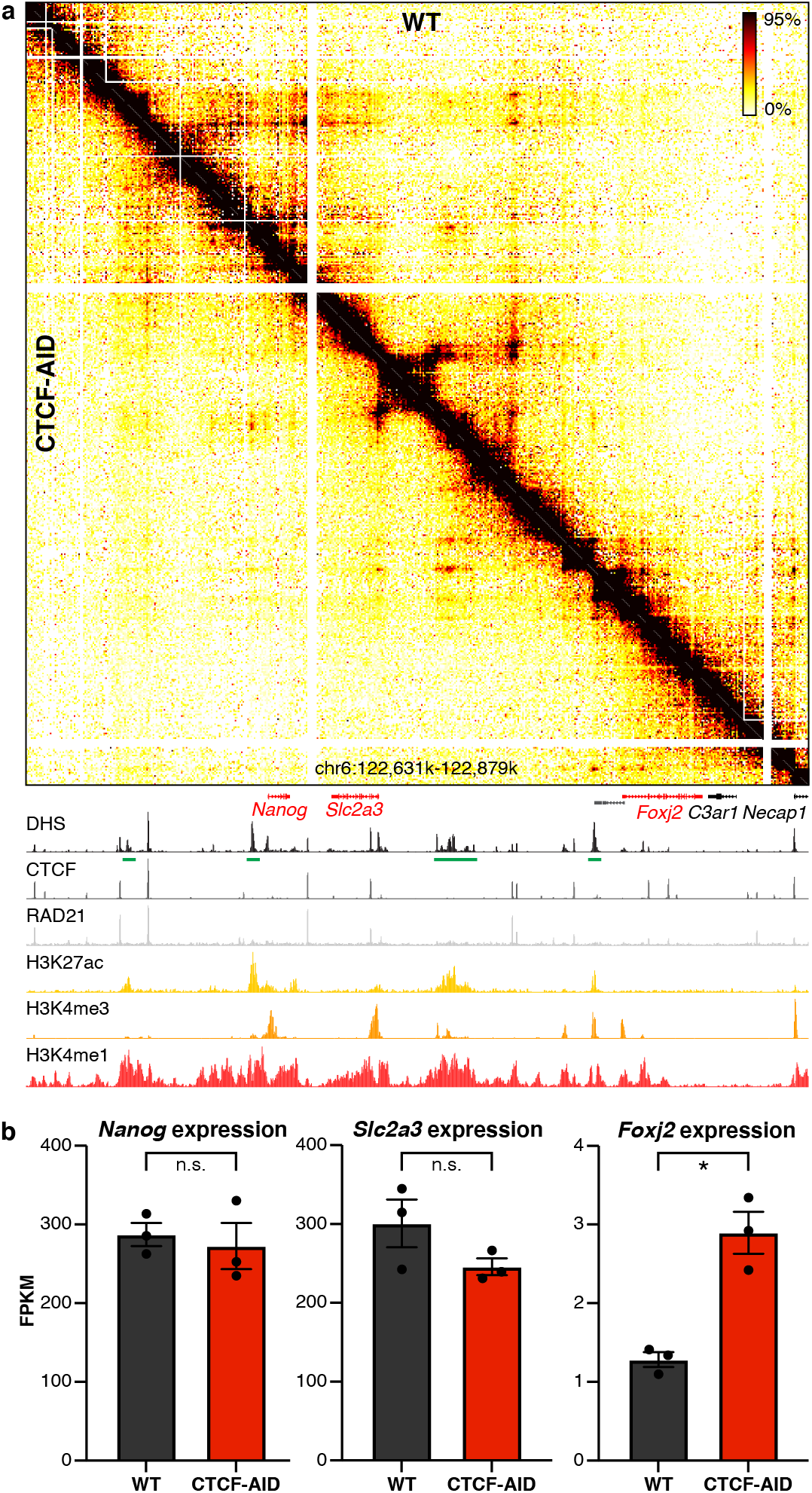
CTCF depletion results in ectopic enhancer-promoter interactions in the *Nanog* locus. **a.** Tiled-MCC contact matrices of the *Nanog* locus in wild type mES cells (top-right) and auxin-treated CTCF-AID mES cells (bottom-left) at 500 bp resolution. Gene annotation (genes of interest in red, coding genes in black, non-coding genes in grey), DNase hypersensitive sites (DHS) and ChIP-seq data for CTCF, Cohesin (RAD21), H3K27ac, H3K4me3 and H3K4me1 are shown below the matrices. The axes of the DHS and ChIP-seq profiles are scaled to signal and have the following ranges: DHS = 0-10.25; CTCF = 0-3092; RAD21 = 0-3414; H3K27ac = 0-58; H3K4me3 = 0-90; H3K4me1 = 0-2064. Enhancers of interest are indicated in green below the DHS profile. **b.** Expression of *Nanog*, *Slc2a3*, and *Foxj2* in untreated (left) and auxin-treated (right) CTCF-AID mES cells, derived from RNA-seq data, normalized for fragments per kilobase of transcript, per million mapped reads (FPKM). The bars represent the average of n=3 replicates and the error bars indicate the standard error of the mean. Significant (*) and non-significant (n.s.) changes in expression are indicated. *Nanog*: P = 0.7030; *Slc2a3*: P = 0.1774; *Foxj2*: P = 0.0001.

To examine the impact of cohesin and CTCF depletion on *cis-*regulatory elements in further detail, we analyzed the effects on local micro-topologies (Figure 4, Supplementary Figure 7). As expected, the pattern of phased nucleosomes, stripes and insulation at CTCF-binding elements are lost upon CTCF depletion. However, surprisingly, the localized insulating properties are not altered by depletion of cohesin, despite complete loss of long-range contacts between CTCF-binding sites upon cohesin depletion (Figure 4, Supplementary Figures 7 and 8). The nano-scale structures of promoters and super-enhancers are also not affected by cohesin depletion, but insulation at CTCF-binding sites within super-enhancers or promoter regions is lost upon CTCF depletion (Figure 4, Supplementary Figure 7). This suggests that CTCF mediates local insulation independent of cohesin-mediated loop extrusion, possibly via recruitment of other factors or its effects on the organization of nucleosomes.

**Figure 4.**
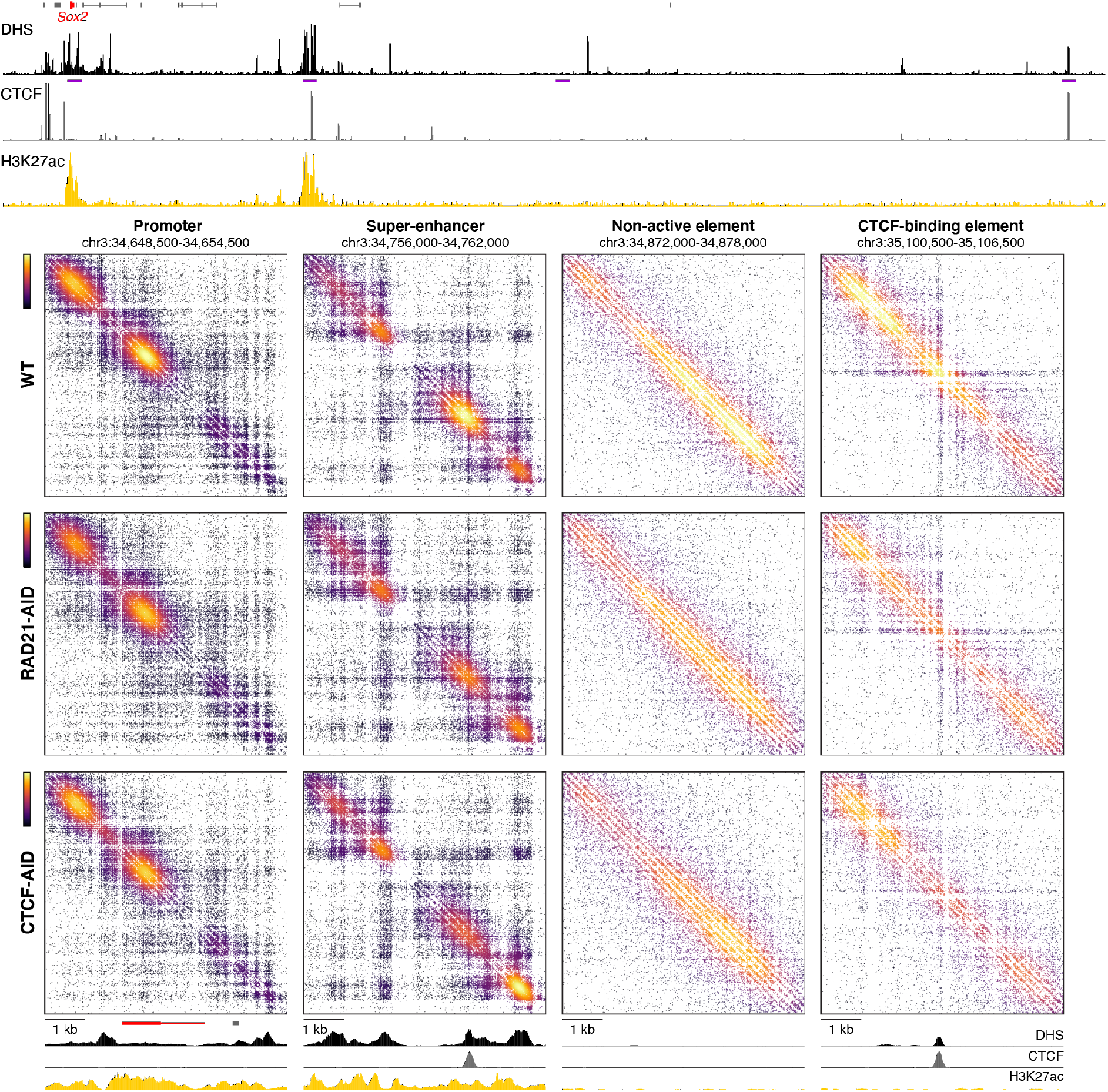
Density plots of ligation junctions identified by Tiled-MCC reveal the micro-topology of *cis*-regulatory elements in the *Sox2* locus and their variable dependence on cohesin and CTCF. Density plots of ligation junctions identified by Tiled-MCC in the *Sox2* locus. Gene annotation (*Sox2* in red, coding genes in black, non-coding genes in grey), DNase hypersensitive sites (DHS), and ChIP-seq data for CTCF and H3K27ac for the extended *Sox2* locus are shown above the plots, with the regions covered in the density plots highlighted in purple. The density plots show a promoter, enhancer, nonactive element and CTCF-binding element in order of the purple highlights and are annotated as described for the panel above. The axes of the DHS and ChIP-seq profiles for CTCF and H3K27ac are fixed and have the following ranges: DHS = 0-5; CTCF = 0-1500; H3K27ac = 0-50.

These observations are confirmed by micro-topology analysis of a larger region at the *Nanog* locus, which contains a promoter, an enhancer and both a strong and a weak CTCF-binding site (Figure 5). Upon cohesin depletion, the specific interactions between the CTCF-binding sites are lost, but the stripe pattern at the strong CTCF-binding site remains, whereas CTCF depletion results in loss of both features. In contrast, the local structures at the enhancer and promoter are not dependent on cohesin and CTCF.

**Figure 5.**
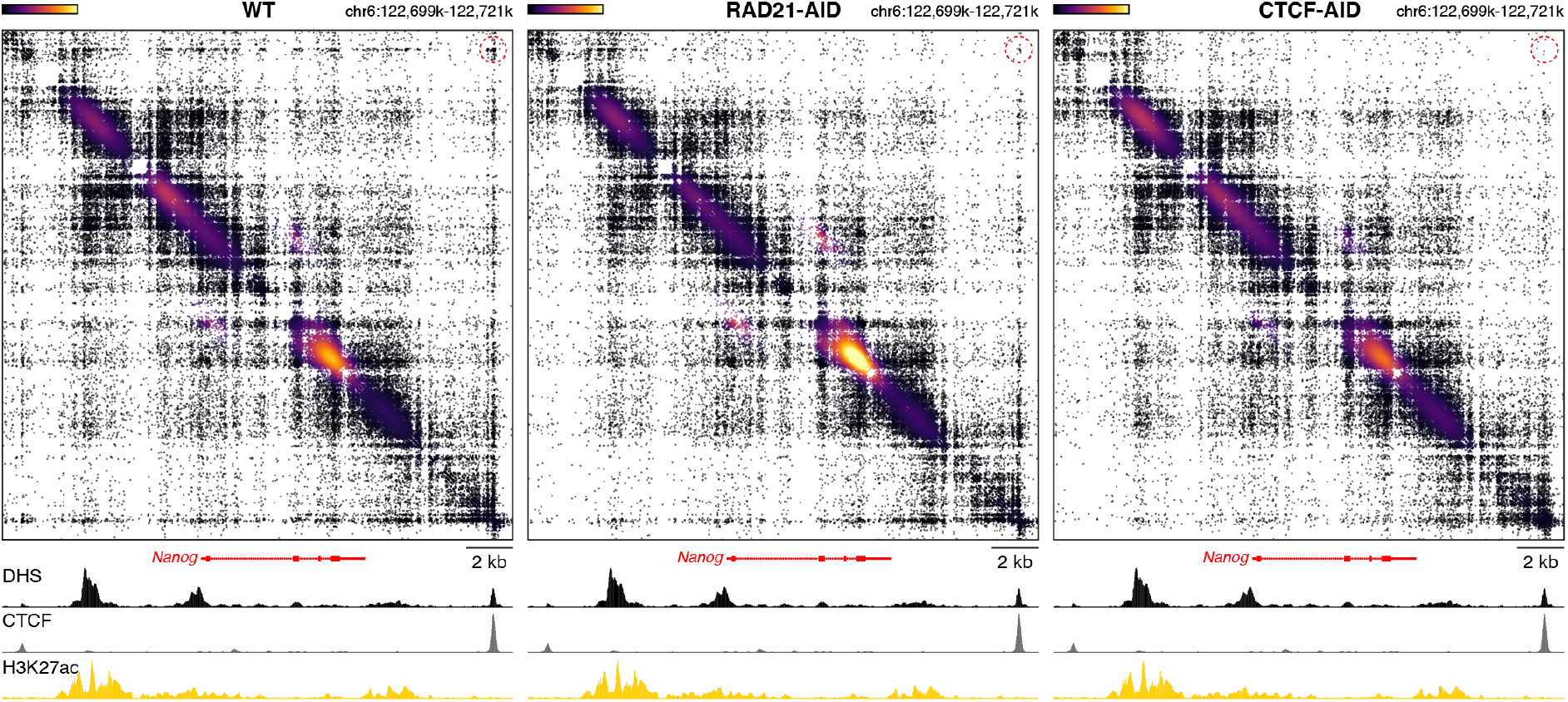
The micro-topology of the *Nanog* locus upon cohesin and CTCF depletion. Density plots of ligation junctions identified by Tiled-MCC in the *Nanog* locus. Gene annotation, DNase hypersensitive sites (DHS), and ChIP-seq data for CTCF and H3K27ac are shown below the plots. The axes of the DHS and ChIP-seq profiles for CTCF and H3K27ac are fixed and have the following ranges: DHS = 0-8; CTCF =0-2050; H3K27ac = 0-60. The interactions between the CTCF-binding elements are indicated with a red circle.

Together, our data show that *cis-*regulatory elements are characterized by distinct internal nano-scale structures, which are independent of cohesin. Interestingly, this includes localized insulation mediated by CTCF-binding sites. However, long-range contacts between CTCF-binding elements and TAD insulation are largely abrogated by cohesin depletion. In contrast, interactions between enhancer and promoters are still present in the absence of cohesin. This is consistent with the findings of a recent preprint^39^. However, our data show that enhancer-promoter interactions are reduced upon cohesin depletion, which indicates that cohesin does contribute to the robustness of enhancer-promoter interactions. This could be mediated by an increase in their interaction frequency through the process of loop extrusion or result from direct bridging of enhancers and promoters by cohesin. These findings are consistent with a recently reported role of the cohesin release factor WAPL in regulating gene expression^40^ and a potential role of cohesin movement along the chromatin in directing regulatory factors, as suggested in a recent preprint^41^. Of interest, it has been suggested that cohesin might be particularly important for mediating very long-range interactions between enhancers and promoters^24,42^. While our data show little change in enhancer-promoter interactions across very small distances (<5 kb), we see a decrease in enhancer-promoter interactions at the relatively small *Pou5f1* and *Nanog* loci, across distances of ~20 and ~50kb, respectively. Furthermore, consistent with the literature, our data show that the CTCF boundaries of TADs contribute to the specificity of enhancer-promoter interactions. Importantly, both the rewiring of enhancer-promoter interactions upon CTCF depletion and the reduced enhancer-promoter interactions upon cohesin depletion are associated with changes in gene expression.

In our study, we have focused on gene loci that are specifically expressed in mES cells. We find consistent reductions in enhancer-promoter interactions and gene expression upon cohesin depletion in all investigated loci. This appears to be in contrast with reported mild effects on gene expression upon loss of cohesin or NIPBL^17,18^. However, our data suggest that cohesin is particularly important for mediating robustness of enhancer-promoter interactions. It is therefore perhaps not surprising that only a subset of genes is mis-regulated upon cohesin depletion, since loss of cohesin might only effect genes whose expression is critically dependent on the robust formation of tissue-specific enhancer-promoter interactions. This is consistent with the reported importance of cohesin for regulating gene expression upon stimuli and during differentiation^43,44^.

Taken together, our high-resolution data identify the existence of complex nano-scale structures within *cis*-regulatory elements, which are independent of loop extrusion. Furthermore, we show that loop extrusion is not essential for the formation of longer-range enhancer-promoter interactions, but an important mechanism to regulate the robustness and specificity of regulatory interactions.

## METHODS

### Cell culture

Wild type (E14), RAD21-mAID-eGFP^21^, and CTCF-AID-eGFP^19^ mouse embryonic stem (mES) cells were cultured on plates pre-coated with 0.1% gelatin (Sigma-Aldrich, G1393) in Glasgow Modified Essential Medium (Gibco, 11710035) supplemented with 10% Fetal Bovine Serum (Gibco, 10270106), 0.01 mM 2-Mercaptoethanol (Gibco, 31350010), 2.4 mM L-glutamine (Gibco, 25030024), 1X nonessential amino acids (Gibco, 11140050), 1X sodium pyruvate (Gibco, 11360070) and 20 ng/ml recombinant mouse Leukemia-Inhibitory Factor (Cell Guidance Systems, GFM200). Cells were grown at 37 °C and 5% CO_2_. Half of the media was exchanged daily and cells were passaged every 2 days by trypsinization (Gibco, 25300054).

### Depletion of RAD21 and CTCF

RAD21-mAID-eGFP and CTCF-AID-eGFP mES cells were passaged and plated 2 days before auxin treatment. On the treatment day, cells were first washed once with phosphate-buffered saline (PBS) and supplemented with equilibrated (37 °C and 5% CO_2_) medium containing 500 μM indole-3-acetic acid (Sigma-Aldrich, I5148) freshly prepared before use. RAD21-mAID-eGFP cells were treated with auxin for 6 hr. Since complete depletion of CTCF takes longer to achieve^19,45^, CTCF-AID-eGFP cells were treated for 48 hr, with exchanging auxin-containing media after the first 24 hr. Alkaline phosphatase testing for the assessment of pluripotency before and after auxin treatment was performed using the Alkaline Phosphatase Detection Kit (EMD Millipore, SCR004).

### Immunoblotting

RAD21-mAID-eGFP and CTCF-AID-eGFP mES cells were grown in 25 cm^2^ flasks until they reached ~80% confluency. Cells were then treated with auxin following the aforementioned treatment conditions for each cell line. Whole-cell extracts were prepared by first dissociating cells by trypsinization, resuspending them in media, pelleting and washing once with PBS. The pellet was then resuspended in 5X the pellet volume in radioimmunoprecipitation assay buffer (RIPA; Thermo Scientific, 89900) with protease inhibitor cocktail (Leupeptin, Carl Roth, CN33.4; Pepstatin A, Carl Roth, 2936.3; PMSF, Carl Roth, 6367.3; Benzamidine hydrochloride, Acros Organics, 105245000, resuspended in ethanol) and 250 U/μL benzonase (Sigma-Aldrich, E1014) and rotated for 1 hr at 4 °C. After centrifugation at maximum speed, whole-cell extract in the supernatant was measured using the Bio-Rad Protein Assay (Bio-Rad, 5000006). 15 μg of protein extracts were mixed with NuPAGE LDS Sample Buffer (Invitrogen, NP0007), supplemented with 50 mM DTT (Carl Roth, 6908.3), and loaded on a NuPAGE 4%-12% Bis-Tris gel (Invitrogen, NP0321PK2). Proteins were transferred to a PVDF membrane using the XCell Blot Module (Invitrogen, EI9051) for 1 hr at 30 V. Membrane blocking was performed in 5% blotting grade milk powder (Carl Roth, T145.2) dissolved in PBS-0.05% Tween-20 (PBS-T) for 1 hr at room temperature. After rinsing the membranes with PBS-T, incubation with primary antibodies was performed as follows: the membrane was cut into two pieces, one piece corresponding to the higher protein ladder size for the incubation of RAD21 and CTCF primary antibodies (RAD21: anti-RAD21 antibody, rabbit polyclonal, Abcam, ab154769, 1:1000; CTCF: anti-CTCF antibody, rabbit polyclonal, Abcam, ab70303, 1:1000), and the second piece with lower protein ladder size for the incubation with the loading control (Histone H3: anti-histone H3 antibody (HRP), Abcam, ab21054, 1:5000). All primary antibodies were diluted in 2% PBS-T and incubated at 4 °C overnight. The next morning, membranes were washed 3X with PBS-T for 5-10 minutes each and incubated with secondary antibody (RAD21 and CTCF: Goat Anti-Rabbit IgG H&L (HRP), Abcam ab205718) in PBS-T in a 1:5000 dilution for 1 hr at room temperature, washed again 3X and analyzed on the Intas ChemoCam Imager HR.

### Tiled-MCC – Fixation

The preparation of Micro-Capture-C (MCC) libraries was performed as previously described^36^. 10 million cells per biological replicate were washed once with PBS, dissociated with trypsin and resuspended in 10 ml of culture media. Fixation was performed by incubating cells with formaldehyde to a final concentration of 2% on a roller mixer for 10 minutes at room temperature. The reaction was quenched by adding cold glycine to a final concentration of 130 mM. Samples were centrifuged at 200 rcf for 5 minutes at 4 °C, the pellet was washed once with cold PBS, centrifuged again and resuspended in 1 ml of cold PBS containing 0.005% digitonin (Sigma-Aldrich, D141).

### Tiled-MCC – Library preparation

Crosslinked and permeabilized cells were centrifuged at 300 rcf for 5 minutes. The supernatant was carefully removed without disrupting the pellet, which was subsequently resuspended in 900 μl nuclease-free water. Cells were then split equally into 3 digestion reactions, such that each reaction contained 3-4 million cells. Titration of different MNase concentrations (NEB, M0247) was performed for each aliquot with MNase concentrations ranging from 30–60 Kunitz U in a total reaction volume of 800 μl containing low-calcium MNase buffer (50 mM Tris-HCl pH 7.5, 1 mM CaCl2). The reaction was then incubated for 1 hr at 37 °C on an Eppendorf Thermomixer at 550 rpm, after which it was quenched with 5 mM of ethylene glycol-bis(2-aminoethylether)-N,N,N’,N’-tetraacetic acid (EGTA, Sigma, E3889). The quenched reaction was subsequently centrifuged for 5 minutes at 300 rcf and the supernatant was carefully discarded. The pellet was resuspended in 1 ml of PBS with 5 mM EGTA. 200 μl was removed as a control for MNase digestion, from which DNA was extracted using the DNeasy blood and tissue kit (Qiagen, 69504) and digestion efficiency was assessed using the Agilent D1000 TapeStation (Agilent Technologies, 5067-5582). MNase reactions that yielded ~180 bp fragments, corresponding to mono-nucleosomes with linkers, were taken further for the subsequent reactions.

To minimize DNA loss, DNA end repair, phosphorylation and ligation were performed in a single tube. However, by controlling the temperature at which the respective enzymes are active, end repair and phosphorylation were performed before ligation. The remaining 800 μl of MNase-digested chromatin resuspended in PBS and EGTA was centrifuged for 5 minutes at 300 rcf. The supernatant was carefully discarded and the pellet was resuspended in DNA ligase buffer (Thermo Scientific, B69) supplemented with dNTPs (NEB, N0447L) at a final concentration of 400 μM each and 2.5 mM EGTA. T4 Polynucleotide Kinase (NEB, M0201L), DNA Polymerase I Large Fragment (Klenow, NEB, M0210L), and T4 DNA ligase (Thermo Scientific, EL0013) were added to final concentrations of 200 U/ml, 100 U/ml, and 300 U/ml, respectively. The reaction was incubated on an Eppendorf Thermomixer at 550 rpm for 2 hr at 37 °C, followed by a 16-hour incubation at 20 °C. Decrosslinking of chromatin was performed using proteinase K (included in the Qiagen DNeasy blood and tissue kit) at 65 °C for > 4 hr, which was followed by DNA extraction using the Qiagen DNeasy blood and tissue kit. The ligation product, referred to as 3C library hereafter, was assessed using the Agilent D1000 TapeStation. A successful ligation is indicated by a significant increase in the fragment size > ~370 bp.

### Tiled-MCC – Sonication and ligation of indexed sequencing adapters

Sonication of 3C libraries was performed using 3-5 μg per library on a Covaris S220 Focused Ultrasonicator with the following conditions: 250-300s: duty cycle 10%; intensity 5; cycles per burst 200, to yield an average fragment size of 200 bp. The sonication quality was assessed using the Agilent D1000 TapeStation. The DNA was purified using Ampure XP beads (Beckman Coulter, A63881). To maximize library complexity, the addition of sequencing adapters was parallelized in triplicate reactions such that each reaction contained 1-2 μg of sonicated 3C library. NEB Ultra II (NEB, 7645S) reagents were used following the manufacturer’s protocol with the following deviations: (1) 2-3 X the amount of adapters was used; (2) all Ampure XP bead clean-up reactions were performed with a DNA sample:bead ratio of 1:1.5; (3) to maximize library complexity and yield, the PCR was performed in triplicate per ligation reaction using the Herculase II PCR reagents (Agilent Technologies, 600677). The parallel library preparations and PCR reactions were subsequently pooled for each reaction.

### Tiled-MCC – Capture oligonucleotide design

Tiled-MCC capture panels were designed to densely cover regions of interest, with oligonucleotides that are 70 nucleotides in length and have an overlap of 35 nucleotides. The sequences were designed and filtered for repetitive sequences using a python-based oligo tool^46^ (https://oligo.readthedocs.io/en/latest/). The panels of double-stranded capture oligonucleotides were ordered from Twist Bioscience (Custom probes for NGS target enrichment).

### Tiled-MCC – Enrichment

The enrichment procedure was performed using the Twist Hybridization and Wash Kit (Twist Bioscience, 101025), Twist Universal Blockers (Twist Bioscience, 100578), and Twist Binding and Purification Beads (Twist Bioscience, 100983). Per hybridization reaction, up to 8 amplified and indexed libraries were multiplexed to a final amount of 1.5 μg in a single 0.2-ml PCR strip-tube. Library pools were dried completely in a vacuum concentrator at 45 °C. Dried DNA was resuspended in 5 μg of mouse Cot-1 DNA and 7 μl of Twist Universal Blockers. In a separate PCR 0.2-ml strip-tube, the probe solution was prepared by mixing 20 μl of Twist Hybridization Mix with 1-2 μl of oligonucleotides and the final volume was adjusted with nuclease-free water. To prepare both the probe solution and the resuspended indexed library pool for hybridization, the probe solution was heated to 95 °C for 2 minutes in a PCR thermal cycler with the lid at 105 °C, then immediately cooled down on ice for 5 minutes. While the probe solution was cooling down on ice, the library pool was heated following the same conditions for 5 minutes. After equilibrating both mixtures to room temperature for 1-2 minutes, the probe solution was added to the library pool, and 30 μl of Twist Hybridization Enhancer was added last. The capture reaction was incubated at 70 °C for 16 hr in a PCR thermal cycler with the lid heated to 85 °C. The hybridization reaction was subsequently mixed with Streptavidin Binding Beads for 30 minutes at room temperature on a shaker. Washing with Twist Wash Buffers 1, 2 and 3 was performed following the Twist Target Enrichment Protocol. Post-hybridization PCR was performed with 11-12 amplification cycles. Enriched library was purified with pre-equilibrated Twist DNA Purification Beads at a ratio of 1:1.8 DNA to beads. DNA quantification and QC validation were performed using the Qubit dsDNA Broad Range Quantification Assay (Life Technologies, Q32850) and the Agilent Bioanalyzer Broad Range DNA kit (Agilent Technologies, 5067-5582), respectively.

### Tiled-MCC – Sequencing

Libraries were sequenced using the Illumina NovaSeq and NextSeq 550 platforms with 150 bp paired-end reads. Depending on the quality of the MCC libraries, sequencing 40-200 million reads per enriched Mb per pooled sample is sufficient for data at 500 bp resolution.

### Tiled-MCC – Analysis

Tiled-MCC analysis was performed using the MCC pipeline^36^ (https://github.com/jojdavies/Micro-Capture-C). Briefly, adapter sequences were removed using Trim Galore (Babraham Institute, v.0.3.1) and paired-end reads were reconstructed using FLASH^47^ (v.1.2.11). Reads were then mapped to the DNA sequences corresponding to the enriched regions of interest with the non-stringent aligner BLAT^48^ (v.35), using a custom-made file containing the sequences of the regions of interest as reference. Based on the mapping by BLAT, the reads in the FASTQ files were split into two or more reads corresponding to the chimeric fragments, and into different files based on the region they mapped to, using the MCCsplitter.pl script. Uninformative reads that did not contain a sequence that mapped to any of the enriched regions of interest were discarded. The split reads in the FASTQ files were subsequently mapped to the mm10 reference genome using Bowtie2^49^ (v.2.3.5). The aligned reads were further processed using the MCCanalyser.pl script. PCR duplicates were removed based on the sonicated ends and ligation junction and allowing for a wobble of ±2 bp. Unique ligation junctions were identified if the fragment ends were less than 5 bp apart in the original read and separated by mapping with BLAT and Bowtie2.

To generate contact matrices, the unique ligation junctions were converted into raw matrices, which were balanced using ICE normalization^50^. To generate density plots, the unique ligation junctions were filtered for a minimum distance ≥ 10 bp and plotted in python as a scatter plot with a color code defined by the local density of the data points.

The data presented in the manuscript represent the averages of 9 replicates for WT samples, 4 replicates for untreated RAD21-AID samples, 6 replicates for auxin-treated RAD21-AID samples, and 6 replicates for auxin-treated CTCF-AID samples. For direct comparisons between Tiled-MCC matrices or density plots, the data were down-sampled to the lowest number of filtered, unique ligation junctions per condition or replicate. The RAD21-AID and CTCF-AID cell lines are derived from the same strain as the WT cells and contact matrices from the untreated RAD21-AID samples are nearly identical to the WT samples (Supplementary Figure 3b). The contact matrices presented in the manuscript therefore show comparisons between auxin-treated RAD21-AID and CTCF-AID samples and WT samples.

### Micro-C analysis

The Micro-C data were downloaded from GSE130275^34^ as files listing valid pairs in the mm10 reference genome, which were converted into ICE-normalized^50^ contact matrices using HiC-Pro^51^.

### ChIP-seq analysis

Alignment to the mm10 reference genome and processing of ChIP-seq data was performed using the NGseqBasic pipeline^52^. ChIP-seq data for CTCF, RAD21, and H3K4me1 were downloaded from GSE30203^53^, GSE94452^22^ and GSE27844^54^, respectively. ChIP-seq data for H3K27ac and H3K4me3 and DNase I hypersensitivity data for mES cells were accessed via ENCODE^55^.

### RNA-seq analysis

The normalized read counts, P-values, and significance scores for genes of interest in RAD21-AID^21^ and CTCF-AID^19^ mES cells were extracted from the original processed data files shared by the authors of the respective articles.

## ACKNOWLEDGEMENTS

A.M.O. is funded by the Max Planck Society. J.O.J.D. is funded by an MRC Clinician Scientist Award (MRC Clinician Scientist Fellowship MR/R008108). We would like to thank Angelika Feldmann, James Rhodes and Rob Klose for providing the RAD21-AID mES cell line and Elphège Nora and Benoit Bruneau for providing the CTCF-AID mES cell line. We are grateful to Patrick Cramer for advice and infrastructure support, Michael Lidschreiber for assistance with bioinformatic analysis, Taras Velychko for support with experiments, and Kerstin Maier and Petra Rus for help with sequencing. We would like to thank Douglas Higgs, Jim Hughes, Damien Downes and Elisa Oberbeckmann for advice and feedback on the manuscript.

## DATA AND CODE AVAILABILITY

Sequencing data have been submitted to the NCBI Gene Expression Omnibus under accession code GSE181694. Scripts for Tiled-MCC analysis are available for academic use through the Oxford University Innovation software store (https://process.innovation.ox.ac.uk/software/p/16529a/micro-capture-c-academic/1).

