## Supplementary Figures for "Analysis of sub-kilobase chromatin topology reveals nano-scale regulatory interactions with variable dependence on cohesin and CTCF"

**Supplementary Figure 1. Tiled-MCC generates highly reproducible contact matrices.**

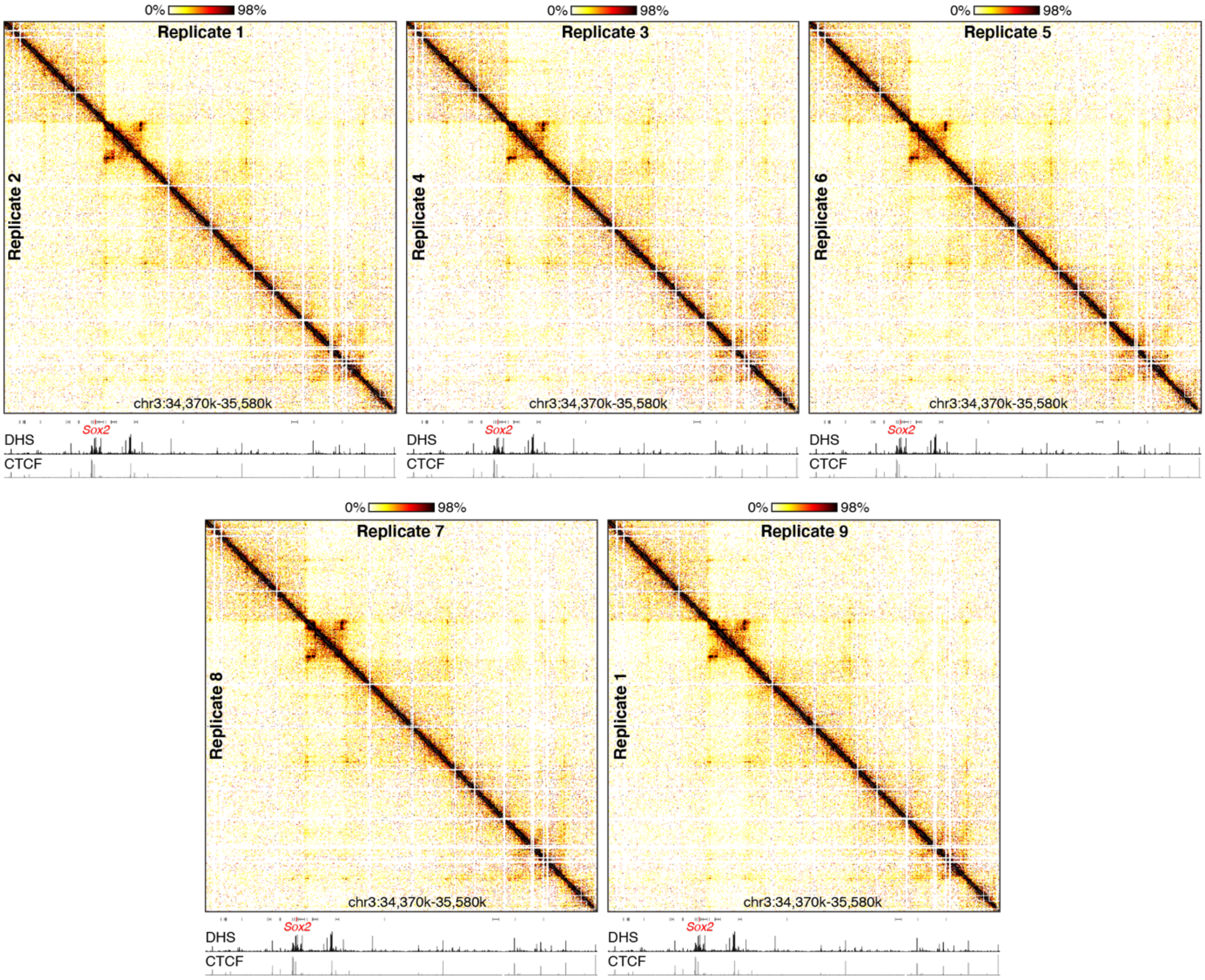


Comparison of Tiled-MCC contact matrices of the *Sox2* locus at 2 kb resolution in 9 independent replicates of wildtype mES cells. Gene annotation (*Sox2* in red, coding genes in black, non-coding genes in grey), DNase hypersensitive sites (DHS) and ChIP-seq data for CTCF are shown below the matrices. The axes of the DHS and ChIP-seq profiles are scaled to signal and have the following ranges: DHS = 0-4.46; CTCF = 0-1833.

**Supplementary Figure 2. Comparison of Tiled-MCC density plots to high-resolution ICE-normalized contact matrices.**

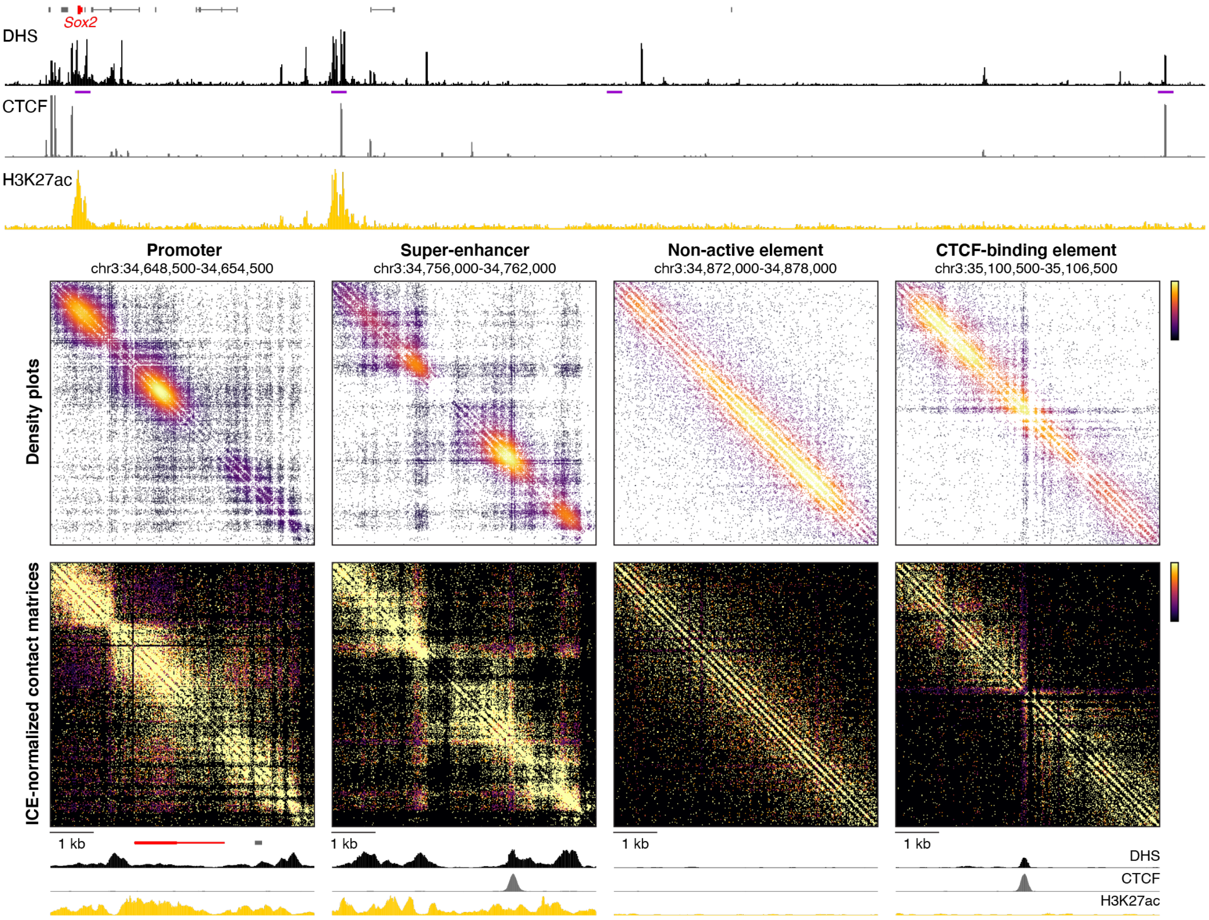


The Tiled-MCC density plots display all identified ligation junctions in regions of interest with a color code defined by the density of the data points. To correct for potential biases resulting from differences in digestion, ligation and capture efficiencies, we compared the density plots to ICE-normalized contact matrices with a resolution of 20 bp, which show very similar patterns of localized interactions.

**Supplementary Figure 3. Quality control of Tiled-MCC data in RAD21-AID mES cells.**

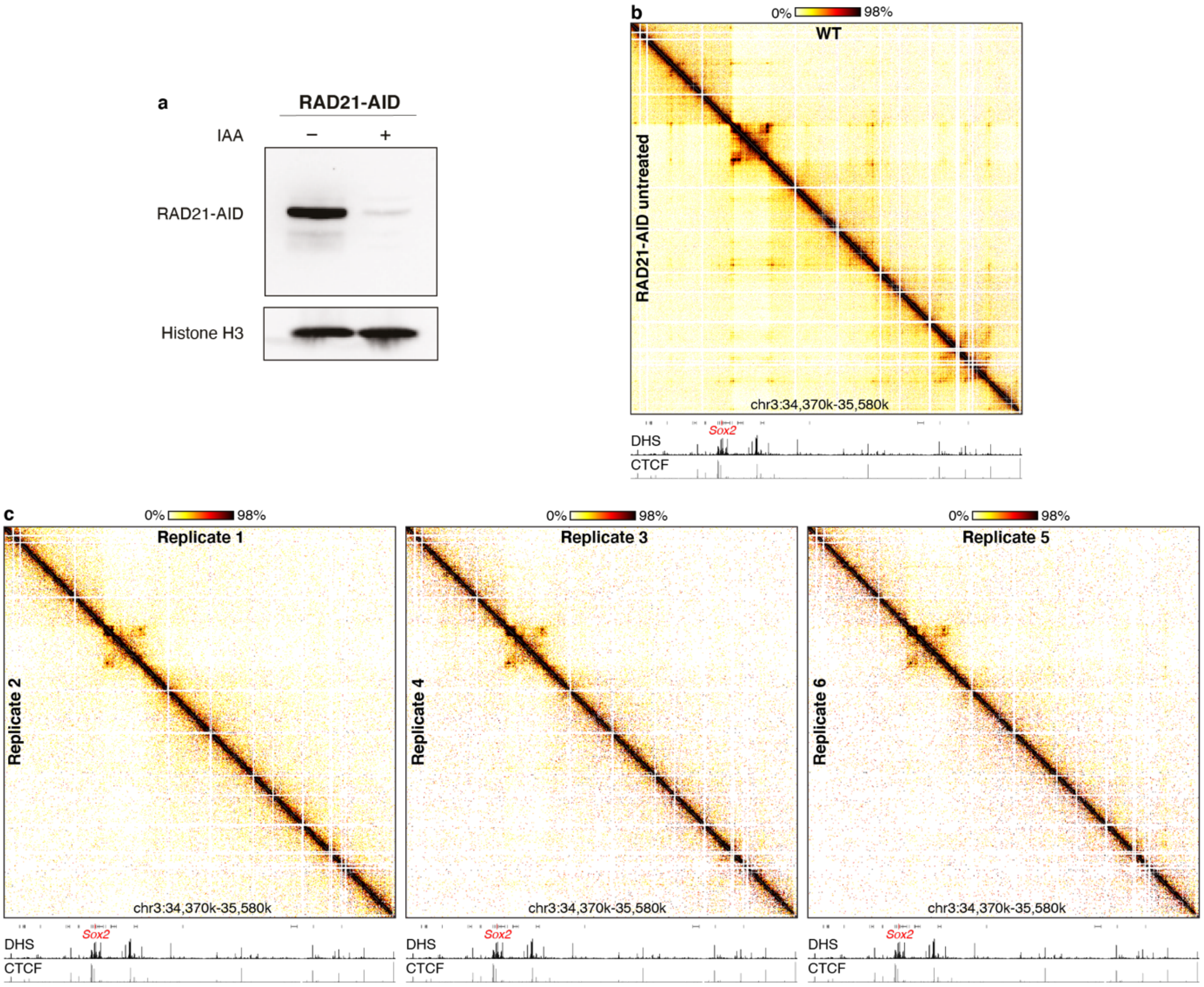


**a.** A representative immunoblot blot for RAD21 in the RAD21-AID mES cell line without treatment and after treatment with indole-3-acetic acid (IAA; auxin) for 6 hours. Histone H3 is shown as a loading control. **b.** Comparison of Tiled-MCC contact matrices of the *Sox2* locus at 1 kb resolution in wildtype (WT) mES cells and untreated RAD21-AID mES cells. Gene annotation (*Sox2* in red, coding genes in black, non-coding genes in grey), DNase hypersensitive sites (DHS) and ChIP-seq data for CTCF are shown below the matrices. The axes of the DHS and ChIP-seq profiles are scaled to signal and have the following ranges: DHS = 0-4.46; CTCF = 0-1833. **c.** Comparison of Tiled-MCC contact matrices of the *Sox2* locus at 2 kb resolution in 6 independent replicates of auxin-treated RAD21-AID mES cells. Annotation below the matrix is the same as in panel b.

**Supplementary Figure 4. Cohesin depletion results in reduced enhancer-promoter interactions.**

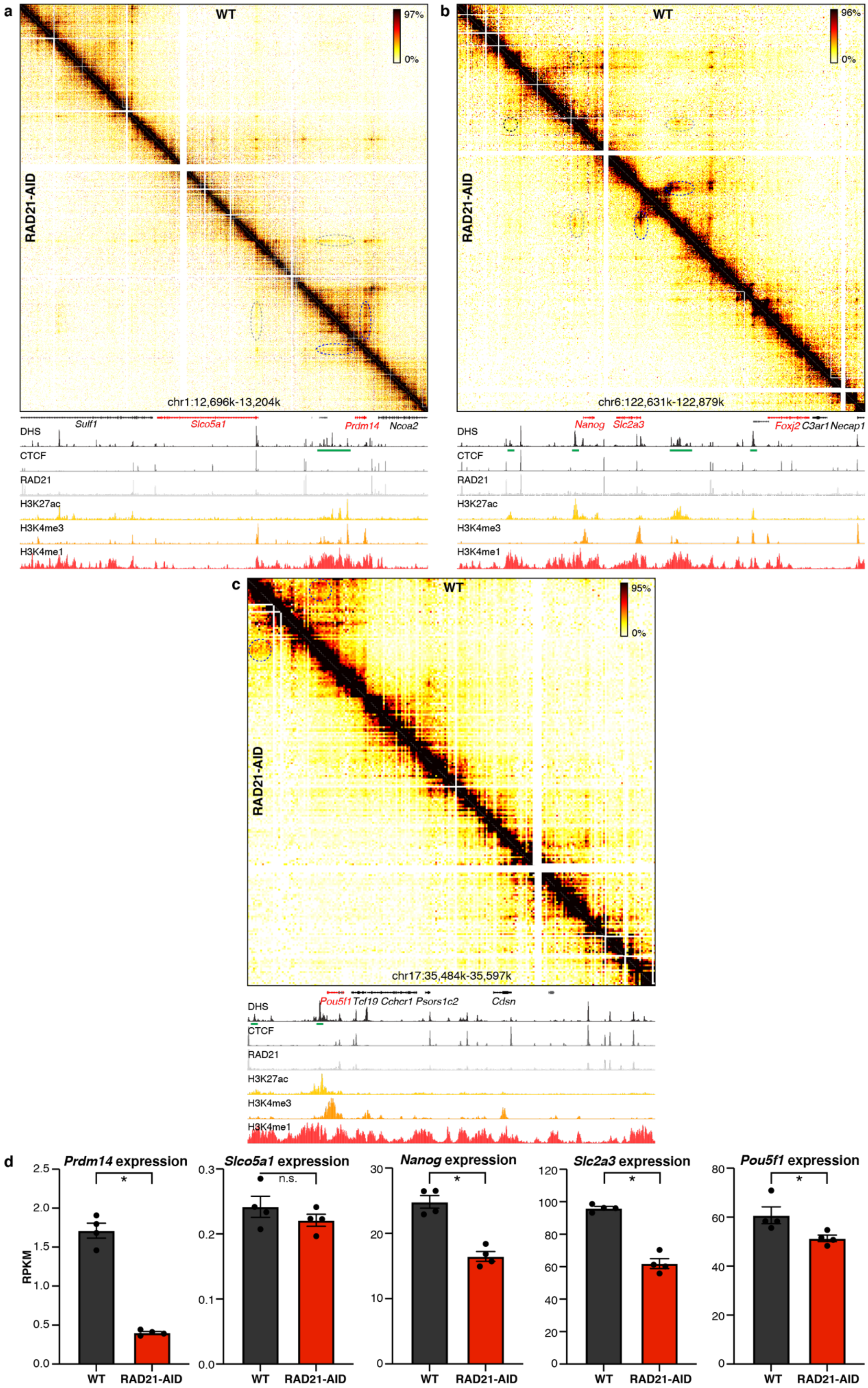

**Supplementary Figure 4. Cohesin depletion results in reduced enhancer-promoter interactions.**

**a-c.** Tiled-MCC contact matrices of the *Prdm14*, *Nanog* and *Pou5f1* loci in wild type (WT) mES cells (top-right) and auxin-treated RAD21-AID mES cells (bottom-left) at 500 bp resolution. Gene annotation (genes of interest in red, coding genes in black, non-coding genes in grey), DNase hypersensitive sites (DHS) and ChIP-seq data for CTCF, Cohesin (RAD21), H3K27ac, H3K4me3 and H3K4me1 are shown below the matrices. The axes of the DHS and ChIP-seq profiles are scaled to signal and have the following ranges; DHS: *Prdm14* = 0-6.45, *Nanog* = 0-10.25, *Pou5f1* = 0-8.65; CTCF: *Prdm14* = 0-2167, *Nanog* = 0-3092, *Pou5f1* = 0-2349; RAD21: *Prdm14* = 0-3032, *Nanog* = 0-3414, *Pou5f1* = 0-4107; H3K27ac: *Prdm14* = 0-44, *Nanog* = 0-58, *Pou5f1* = 0-81; H3K4me3: *Prdm14* = 0-26, *Nanog* = 0-90, *Pou5f1* = 0-64; H3K4me1: *Prdm14* = 0-2170, *Nanog* = 0-2064, *Pou5f1* = 0-1641. Enhancers of interest are indicated in green below the DHS profile. **a.** In WT cells, the enhancers form strong interactions with the *Prdm14* promoter (blue) and weak interactions with the *Slco5a1* promoter across a CTCF boundary (grey). Upon cohesin depletion, the interactions between the enhancers and the *Prdm14* promoter remain present, but are reduced in intensity. **b**. The *Nanog* locus contains several enhancers. Upon depletion of cohesin, the interactions between the *Nanog* promoter and the enhancer far upstream (black) are reduced, whereas its interactions with the enhancer directly upstream and the enhancer downstream (grey) appear unchanged. The *Slc2a3* promoter also interacts with the downstream enhancer (blue); these interactions are weakened but remain present upon cohesin depletion. **c.** *Pou5f1* is regulated by an enhancer directly upstream and an enhancer further upstream (blue). The interactions with the far-upstream enhancer are reduced in intensity upon cohesin depletion. **d.** Expression of *Prdm14*, *Slco5a1*, *Nanog*, *Slc2a3*, and *Pou5f1* in untreated (left) and auxin-treated (right) RAD21-AID mES cells, derived from RNA-seq data, normalized for reads per kilobase of transcript, per million mapped reads (RPKM). The bars represent the average of n=4 replicates and the error bars indicate the standard error of the mean. Significant (*) and non-significant (n.s.) changes in expression are indicated. *Prdm14*: P = 3.64E-45; *Slco5a1*: P = 0.388; *Nanog*: P = 2.76E-14; *Slc2a3*: P = 3.64E-16; *Pou5f1*: P = 0.00586.

**Supplementary Figure 5. Quality control of Tiled-MCC data in CTCF-AID mES cells.**

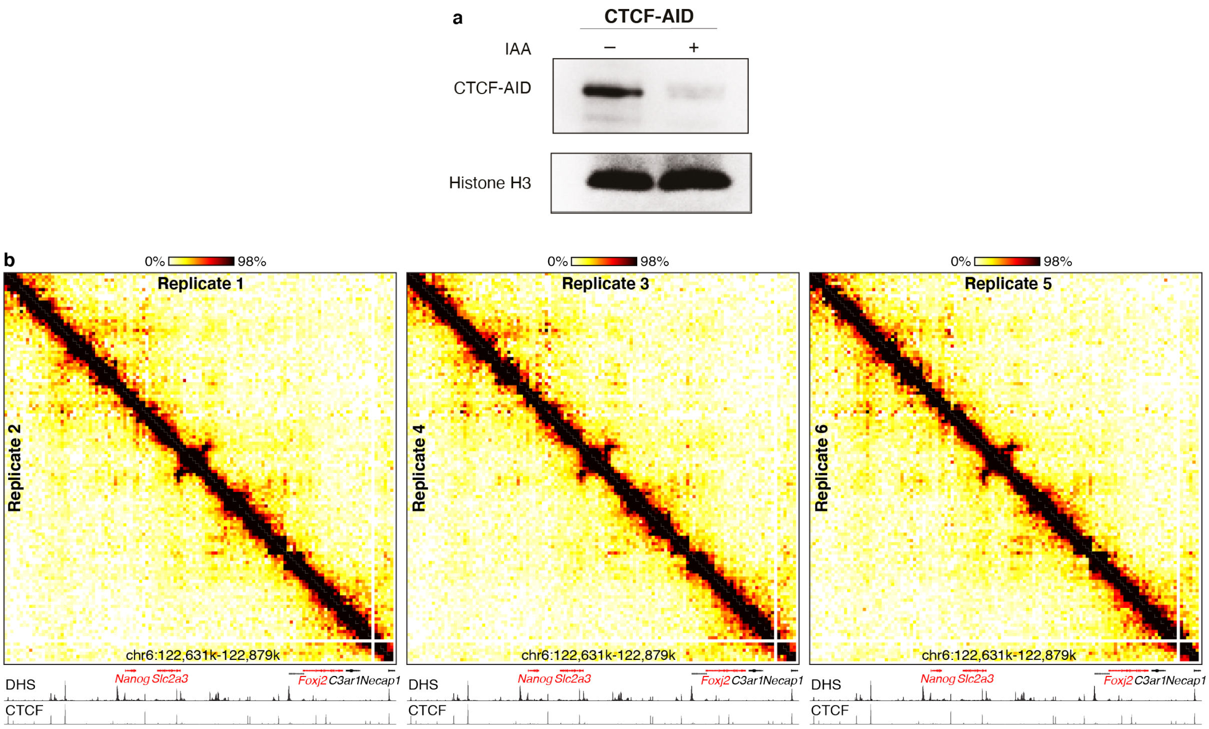


**a.** A representative immunoblot for CTCF in the CTCF-AID mES cell line without treatment and after treatment with indole-3-acetic acid (IAA; auxin) for 48 hours. Histone H3 is shown as a loading control. **b.** Comparison of Tiled-MCC contact matrices of the *Nanog* locus at 2 kb resolution in 6 independent replicates of auxin-treated CTCF-AID mES cells. Gene annotation (genes of interest in red, coding genes in black, non-coding genes in grey), DNase hypersensitive sites (DHS) and ChIP-seq data for CTCF are shown below the matrices. The axes of the DHS and ChIP-seq profiles are scaled to signal and have the following ranges: DHS = 0-10.25; CTCF = 0-3092.

**Supplementary Figure 6. CTCF depletion can result in ectopic enhancer-promoter interactions but does not otherwise mediate the strength of enhancer-promoter interactions.**

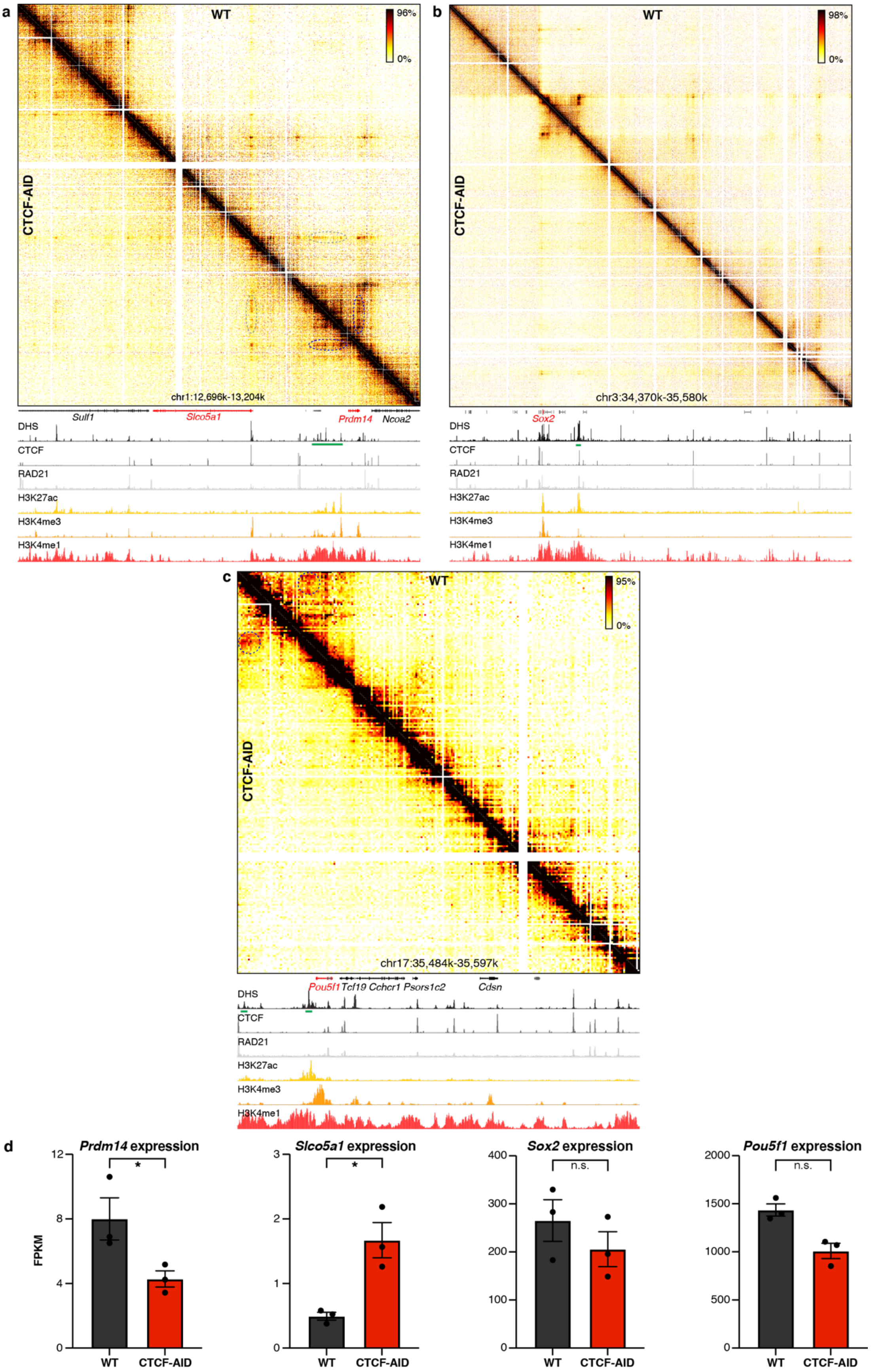

**Supplementary Figure 6. CTCF depletion can result in ectopic enhancer-promoter interactions but does not otherwise mediate the strength of enhancer-promoter interactions.**

**a-c.** Tiled-MCC contact matrices of the *Prdm14*, *Sox2* and *Pou5f1* loci in wild type (WT) mES cells (top-right) and auxin-treated CTCF-AID mES cells (bottom-left) at 500 bp resolution. Gene annotation (genes of interest in red, coding genes in black, non-coding genes in grey), DNase hypersensitive sites (DHS) and ChIP-seq data for CTCF, Cohesin (RAD21), H3K27ac, H3K4me3 and H3K4me1 are shown below the matrices. The axes of the DHS and ChIP-seq profiles are scaled to signal and have the following ranges; DHS: *Prdm14* = 0-6.45, *Sox2* = 0-4.46, *Pou5f1* = 0-8.65; CTCF: *Prdm14* = 0-2167, *Sox2* = 0-1833, *Pou5f1* = 0-2349; RAD21: *Prdm14* = 0-3032, *Sox2* = 0-3318, *Pou5f1* = 0-4107; H3K27ac: *Prdm14* = 0-44, *Sox2* = 0-48, *Pou5f1* = 0-81; H3K4me3: *Prdm14* = 0-26, *Sox2* = 0-82, *Pou5f1* = 0-64; H3K4me1: *Prdm14* = 0-2170, *Sox2* = 0-1826, *Pou5f1* = 0-1641. Enhancers of interest are indicated in green below the DHS profile. **a.** CTCF depletion results in a subtle increase in the contacts between the *Prdm14* enhancers and the upstream *Slco5a1* promoter (grey), whereas interactions between the enhancers and the *Prdm14* promoter (blue) are slightly reduced in intensity. **b.** Depletion of CTCF reduces interactions between CTCF-binding sites in the *Sox2* locus but does not result in robust changes in the enhancer-promoter interactions. **c.** The interactions between *Pou5f1* and its enhancer (blue) do not change substantially upon CTCF deletion. **d.** Expression of *Prdm14*, *Slco5a1*, *Sox2*, and *Pou5f1* in untreated (left) and auxin-treated (right) CTCF-AID mES cells, derived from RNA-seq data, normalized for fragments per kilobase of transcript, per million mapped reads (FPKM). The bars represent the average of n=3 replicates and the error bars indicate the standard error of the mean. Significant (*) and non-significant (n.s.) changes in expression are indicated. *Prdm14*: P = 0.0006; *Slco5a1*: P = 5.00E-05; *Sox2*: P=0.098; *Pou5f1*: P = 0.02515.

**Supplementary Figure 7. Density plots of ligation junctions identified by Tiled-MCC reveal the micro-topology of *cis*-regulatory elements in the *Prdm14* locus and their variable dependence on cohesin and CTCF.**

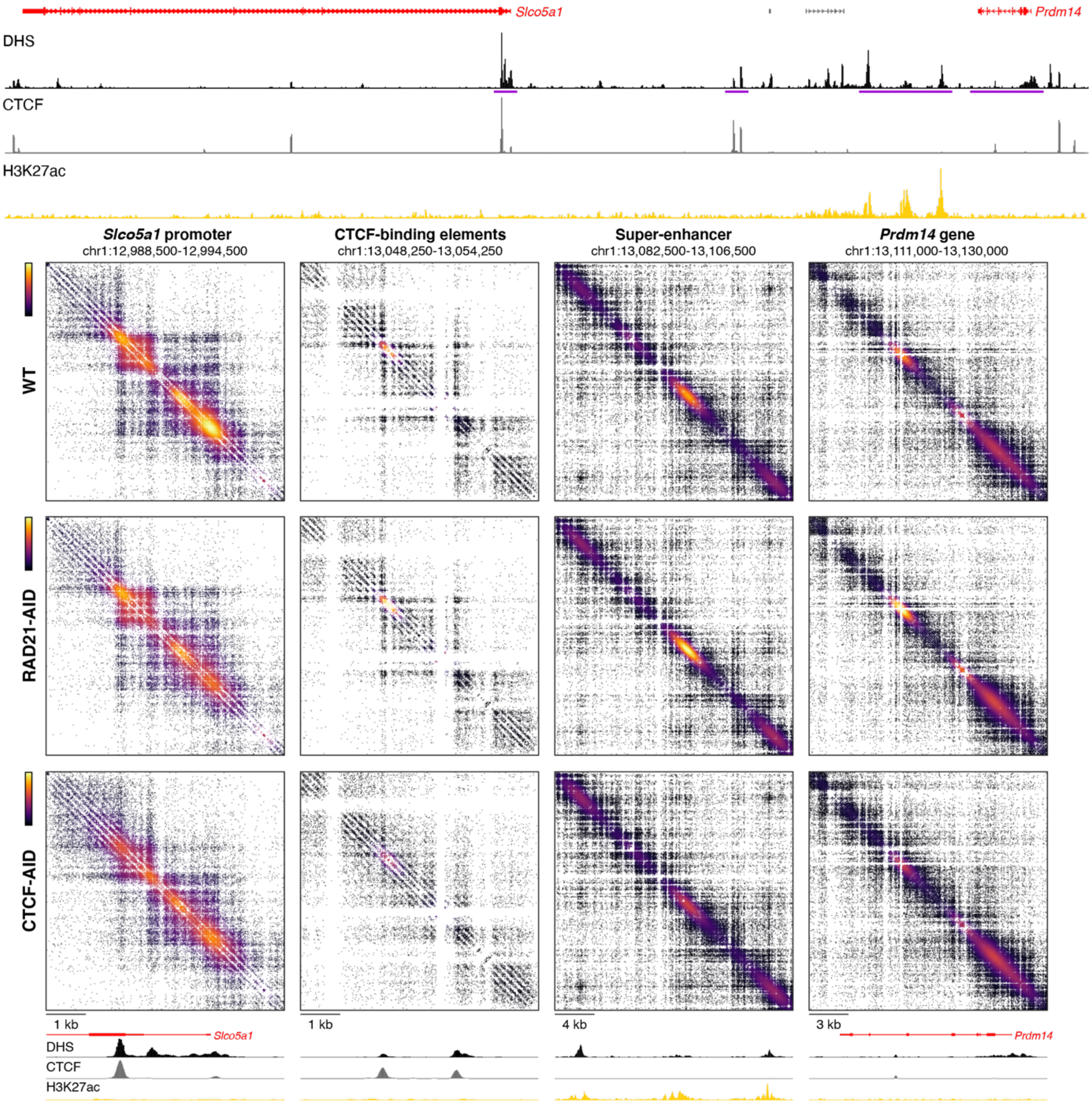


Density plots of ligation junctions identified by Tiled-MCC in the *Prdm14* locus. Gene annotation (genes of interest in red, coding genes in black, non-coding genes in grey), DNase hypersensitive sites (DHS), and ChIP-seq data for CTCF and H3K27ac for the extended *Prdm14* locus are shown above the plots, with the regions covered in the density plots highlighted in purple. The density plots show the *Slco5a1* promoter, CTCF-binding elements, enhancers, and the *Prdm14* gene in order of the purple highlights and are annotated as described for the panel above. The axes of the DHS and ChIP-seq profiles for CTCF and H3K27ac are fixed and have the following ranges: DHS = 0-6.5; CTCF = 0-2200; H3K27ac = 0-50.

**Supplementary Figure 8. The micro-topology of CTCF-binding elements in the *Sox2* locus upon cohesin and CTCF depletion.**

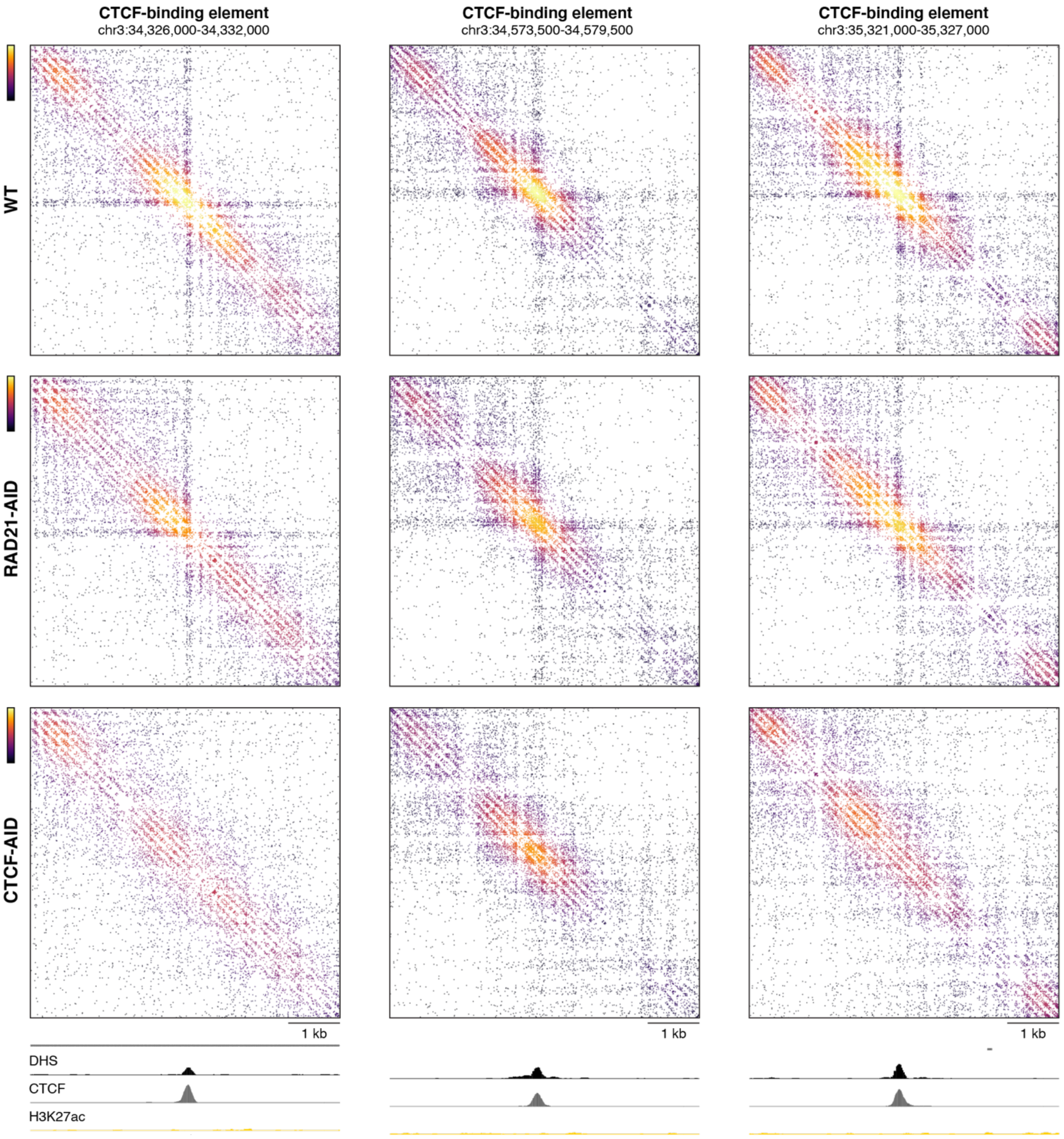


Density plots of ligation junctions identified by Tiled-MCC surrounding CTCF-binding elements in the *Sox2* locus. Gene annotation (non-coding genes in grey), DNase hypersensitive sites (DHS), and ChIP-seq data for CTCF and H3K27ac are shown below the plots. The axes of the DHS and ChIP-seq profiles for CTCF and H3K27ac are fixed and have the following ranges: DHS = 0-5; CTCF = 0-1500; H3K27ac = 0-50.
